# Comprehensive benchmarking of single cell RNA sequencing technologies for characterizing cellular perturbation

**DOI:** 10.1101/2020.11.25.396523

**Authors:** Verboom Karen, Alemu T Assefa, Nurten Yigit, Jasper Anckaert, Niels Vandamme, Dries Rombaut, Yvan Saeys, Olivier Thas, Frank Speleman, Kaat Durinck, Jo Vandesompele

## Abstract

Technological advances in transcriptome sequencing of single cells continues to provide an unprecedented view on tissue composition and cellular heterogeneity. While several studies have compared different single cell RNA-seq methods with respect to data quality and their ability to distinguish cell subpopulations, none of these studies investigated the heterogeneity of the cellular transcriptional response upon a chemical perturbation. In this study, we evaluated the transcriptional response of NGP neuroblastoma cells upon nutlin-3 treatment using the C1, ddSeq and Chromium single cell systems. These devices and library preparation methods are representative for the wide variety of platforms, ranging from microfluid chips to droplet-based systems and from full transcript sequencing to 3-prime end sequencing. In parallel, we used bulk RNA-seq for molecular characterization of the transcriptional response. Two complementary metrics to evaluate performance were applied: the first is the number and identity of differentially expressed genes as defined in consensus by two statistical models, and the second is the enrichment analysis of biological signals. Where relevant, to make the data more comparable, we downsampled sequencing library size, selected cell subpopulations based on specific RNA abundance features, or created pseudobulk samples. While the C1 detects the highest number of genes per cell and better resembles bulk RNA-seq, the Chromium identifies most differentially expressed genes, albeit still substantially fewer than bulk RNA-seq. Gene set enrichment analyses reveals that detection of a limited set of the most abundant genes in single cell RNA-seq experiments is sufficient for molecular phenotyping. Finally, single cell RNA-seq reveals a heterogeneous response of NGP neuroblastoma cells upon nutlin-3 treatment, revealing putative late-responder or resistant cells, both undetected in bulk RNA-seq experiments.

## INTRODUCTION

Almost a decade ago, the first single cell RNA-seq study was published, in which cells were manually isolated and polyadenylated transcripts were captured using oligo(dT) reverse transcription primers (1). Since then, numerous single cell RNA-seq methods and devices have emerged, unveiling an unanticipated cellular heterogeneity that was not very well recognized through classic bulk cell population gene expression profiles. As such, single cell RNA-seq enabled the identification of subtle differences among cells and the detection of rare or novel subpopulations. This has led to revolutionary discoveries in several research fields, including cancer (2, 3) and embryonic development (4–6). The first automated single cell isolation devices used flow cytometry or microfluidic chips and could only capture a hundred cells. Most RNA library preparation protocols for these systems provide full gene body read coverage, enabling mutation and splice isoform analysis on top of classic gene abundance profiling (7–10). Using these methods, single cells can be visualized to remove cell doublets or select cells of interest. Later, commercially available and custom made droplet-based methods, such as Chromium, ddSeq and InDrop, were developed increasing the throughput to thousands of cells and reducing the cost per cell considerably (11–15). One limitation is that these methods typically sequence only the 3’ end of a transcript, reducing the analyses to gene expression profiling. Further, these droplet-based methods typically quantify only the most abundant genes, excluding for instance the detection of medium to low abundant mRNAs and the majority of long non-coding RNAs (lncRNAs). Consequently, lower complexity sequencing libraries are generated using these droplet-based methods, resulting in more PCR bias. Fortunately, this bias can to a large degree be reduced through unique molecular indices (UMI), incorporated in the to-be-sequenced molecules in droplet-based systems (14–17). Also, virtually all these initial single cell RNA-seq methods only capture polyadenylated transcripts, ignoring the vast non-polyadenylated part of the transcriptome. Since flow cytometry and microfluidic chip based methods are mostly open systems, single cell total RNA-seq protocols were recently custom developed enabling the sequencing of both polyadenylated as well as non-polyadenylated transcripts (18–20). The extensive advances in the single cell RNA-seq technologies raises the question which method is best suited for a particular application. While several studies compared single cell RNA-seq methods in terms of data quality, costs, reproducibility, and the ability to discriminate subpopulations, our study focuses on the added value of three single-cell RNA-seq technologies for differential gene expression analysis and assessment of transcriptional heterogeneity (21–25). Therefore, cell cycle synchronized NGP neuroblastoma cells were treated with the TP53 activator nutlin-3, whose transcriptional effects are well-characterized in bulk, resulting in activation of the *TP53* pathway and consequently in cell cycle arrest and apoptosis (26, 27). Single cell RNA-seq of this well-characterized model system has been performed using three commercially available single cell devices, C1 (Fluidigm), ddSeq (Bio-Rad, Illumina) and Chromium (10X Genomics), representing microfluidic chip-based and droplet-based single cell RNA-seq platforms, and a range of throughputs from 96, 300 or more than 10,000 cells per condition, respectively. As a reference, the same experiment was also performed using bulk RNA-seq of ten replicates per group.

Despite the lower number of differentially expressed genes detected in single cell RNA-seq experiments compared to bulk population analysis, the biological signal can be faithfully recognized through gene set enrichment analysis of data from all tested single cell devices. Furthermore, we show that single cell transcriptome analysis reveals a certain degree of cellular heterogeneity in response to nutlin-3 treatment, possibly pinpointing to late-responding or resistant cells, hidden in bulk RNA-seq experiments.

## METHODS

### Cell lines

The neuroblastoma cell line NGP is a kind gift of Prof. R. Versteeg (Amsterdam, the Netherlands). Cells were maintained in RPMI-1640 medium (Life Technologies, 52400-025) supplemented with 10 % fetal bovine serum (PAN Biotech, P30-3306), 1 % of L-glutamine (Life Technologies, 15140-148) and 1 % penicillin/streptomycin (Life Technologies, 15160-047) (referred to as complete medium) at 37 °C in a 5 % CO_2_ atmosphere. Short tandem repeat genotyping was used to validate cell line authenticity prior to performing the described experiments and verification of absence of mycoplasma was done on a monthly basis.

### Cell cycle synchronization and nutlin-3 treatment of NGP cells

NGP cells were synchronized using serum starvation prior to nutlin-3 treatment. First, cells were seeded at low density for 48 hours in complete medium. Then, cells were refreshed with serum-free medium for 24 hours. Finally, the cells were treated with either 8 μM of nutlin-3 (Cayman Chemicals, 10004372, dissolved in ethanol) or vehicle (ethanol). Cells were trypsinized (Gibco, 25300054) 24 hours post treatment and harvested for single cell analysis, bulk RNA isolation and cell cycle analysis.

### Cell cycle analysis

Four million cells were washed with PBS (Gibco, 14190094) and the pellet was resuspended in 300 μl PBS. Next, 700 μl of 70 % ice-cold ethanol was added dropwise while vortexing to fix the cells. Cells were stored at −20 °C for at least 1 hour. After incubation, cells were washed with PBS and the pellet was resuspended in 1 ml PBS containing RNAse A (Qiagen, 19101) at a final concentration of 0.2 mg/ml. After 1 hour incubation at 37 °C, propidium iodide (BD biosciences, 556463) was added to a final concentration of 40 μg/ml. Samples were loaded on a S3 cell sorter (Bio-Rad) and analyzed using the FlowJo v.10 software.

### RNA isolation and cDNA synthesis

Total RNA was isolated using the miRNeasy mini kit (Qiagen, 217084) with DNA digestion on-column according to the manufacturer’s instructions. RNA concentration was measured using spectrophotometry (Nanodrop 1000, Thermo Fisher Scientific). cDNA was synthesized using the iScript Advanced cDNA synthesis kit (Bio-Rad, 1708897) using 500 ng RNA as input in a 20 μl reaction. cDNA was diluted to 2.5 ng/μl with nuclease-free water prior to RT-qPCR measurements.

### Reverse transcription quantitative PCR

PCR mixes containing 2.5 μl 2x SsoAdvansed SYBR qPCR supermix (Bio-Rad, 04887352001), 0.25 μl each forward and reverse primer (5 μM, IDT), and 2 μl diluted cDNA (5 ng total RNA equivalents) were analyzed on the LightCycler 480 instrument (Roche) using two replicates. Expression levels of targets *CDKN1A*, *BAX* and *BBC3* were normalized using four stable reference genes (*SDHA, YWHAZ, TBP, HPRT1*). RT-qPCR data was analyzed using the qbase+ software v3.0 (Biogazelle). Primer sequences are available in Supplementary Table 1.

### Bulk RNA library preparation of NGP cells

The RNA of ten biological replicates of NGP cells treated with either nutlin-3 or vehicle, without serum starvation was extracted using the RNeasy mini kit. The RNA concentration was measured using spectrophotometry (Nanodrop 1000) and quality ascertained using the fragment analyzer (Advanced Analytical). 100 ng of total RNA was used as input for the TruSeq stranded mRNA library prep kit (Illumina, 20020594), according to manufacturer’s instructions.

### Single cell RNA library preparation of C1 isolated NGP cells

Cells were washed with PBS and centrifuged at 300 g for 5 minutes. Pellets of vehicle treated cells were resuspended and incubated in 1 ml pre-warmed (37 °C) cell tracker (CellTracker Green BODIPY Dye, Thermo fisher Scientific, C2102) for 20 minutes at room temperature. After incubation, cells were washed in PBS and resuspended in 1 ml wash buffer (Fluidigm, 100-6201). An equal number of stained (vehicle treated) and non-stained (nutlin-3 treated) cells were mixed and diluted to 300,000 cells per ml. Suspension buffer was added to the cells in a 3:2 ratio and 6 μl of this mix of was loaded on a primed C1 Single-Cell Auto Prep Array for mRNA Seq (Fluidigm, 100-6041) designed for medium-sized cells (10-17 μm). Single cell polyA[+] RNA-seq on the C1 was performed using the SMART-Seq v4 Ultra Low Input RNA Kit for the Fluidigm C1 System (Takara, 635026) according to manufacturer’s instructions. One microliter of the ERCC spike-in mix was diluted in 999 μl loading buffer to get a 1/1000 dilution of the ERCC spikes. One microliter of this dilution was added to the 20 μl lysis mix. The quality of the cDNA was checked for 11 random single cells on the Fragment Analyzer. The concentration of the cells was measured using the quantifluor dsDNA kit (Promega, E2670) and glomax (Promega) according to manufacturer’s instructions. The samples were 1/5 diluted in C1 harvest reagent (Fluidigm). Next, library prep was performed using the Nextera XT library prep kit (Illumina, FC-131-1096) according to manufacturer’s instructions, followed by quality control on the Fragment Analyzer.

### Single cell RNA library preparation of ddSeq isolated NGP cells

Single cell RNA-seq on the ddSeq system (Bio-Rad) was performed using the SureCell WTA 3’ library prep kit (Illumina, 20014279) according to manufacturer’s instructions with minor modifications. Four samples were prepared: (1) nutlin-3 treated cells with ERCC spikes diluted to 1/1000 (N704 index), (2) nutlin-3 treated cells with ERCC spikes diluted to 1/10,000 (N705 index), (3) vehicle treated cells with ERCC spikes diluted to 1/1000 (N706 index) and (4) vehicle treated cells with ERCC spikes diluted to 1/10,000 (N707 index). Cells were diluted to 5000 cells/μl and ERCC spikes were diluted to 1/500 and 1/5000. Cells and ERCC spikes were mixed 1:1 resulting in a final concentration of 2500 cells/μl and a dilution of 1/1000 and 1/10,000 for the ERCC spikes, respectively. After library preparation, the quality of the RNA libraries was confirmed on the Bioanalyzer (Agilent).

### Single cell RNA library preparation of Chromium isolated NGP cells

Single cell RNA-seq on the Chromium system (10X Genomics) was performed for nutlin-3 (SI-GA-8E index) and vehicle (SI-GA-8D index) treated NGP cells using the GemCode Single Cell 3’ Gel Bead and Library Kit (V2 chemistry, 10X Genomics, PN-120237, PN-120236, PN-120262) according to manufacturer’s instructions with minor modifications. Cells were centrifuged at 4 °C at 400 g and resuspended in PBS + 0.04 % BSA to yield an estimated concentration of 1000 cells/μl. 3.5 μl of the cell suspension was used to obtain a cell recovery of about 2000 cells per sample. Per sample, 2.5 μl of an 1/10 dilution of ERCC spikes was added to the mastermix. After library preparation, the quality of the RNA libraries was confirmed on the Bioanalyzer.

### Library sequencing

Bulk RNA-seq libraries were quantified using KAPA library quantification kit (Roche) and diluted to 4 nM. 1.2 pM of the library was paired-end sequenced on a NextSeq 500 (Illumina) with a read length of 75 bp. The C1 RNA-seq libraries were quantified using the KAPA library quantification kit and libraries were diluted to 4 nM. 1.5 pM of the library was single-end sequenced on a NextSeq 500 (Illumina) with a read length of 75 bp. The ddSeq RNA-seq libraries were quantified using the Qubit dsDNA HS kit (Thermo Fischer Scientific, Q32854) and libraries were diluted to 2 nM. 3 pM of the library was paired-end sequenced on a NextSeq 500 with a read length of 68 and 75 bp and a custom sequencing primer included in the SureCell WTA 3’ library prep kit. The Chromium RNA libraries were quantified using the KAPA library quantification kit and libraries were diluted to 4 nM. 1.2 pM of the library was paired-end sequenced twice on a NextSeq 500 with a read length of 26 and 98 bp.

### Data analysis of the bulk RNA sequencing data

Raw fastq files were processed with Kallisto (v.0.43.1) (28) using Ensembl (v.91) annotation (29) .

### Data analysis of the C1 RNA sequencing data

To assess the quality of the data, the reads were mapped using STAR (v.2.5.3) (30) on the hg38 genome including the full ribosomal DNA (45S, 5.8S and 5S) and mitochondrial DNA sequences. The STAR parameters were set to retain only primary mapping reads, meaning that for multi-mapping reads only the best scoring location is retained. Genes were quantified by Kallisto (v.0.43.1) (28) using Ensembl (v.91) (29) annotation supplemented with the ERCC spike-in RNA sequences.

### Data analysis of the ddSeq RNA sequencing data

To analyze the ddSeq data, ddSeeker, a custom pipeline based on the Drop-seq Core Computational Protocol (version 2.0.0 −9/28/18), was used (31). ddSeeker.py was run on paired-end gzipped fastq files with default parameters using Python (v.3.6.4), pysam (v.0.14) and Biopython (v.1.71). First, fastq files were converted to unaligned BAM files using Picard FastqToSam. These BAM files were subsequently tagged with both cell (XC) and molecular (XM) barcodes using TagBamWithReadSequenceExtended. Next, these tagged BAM files were filtered to remove reads below the base quality threshold (XQ) and to remove erroneous barcodes (XE). The SMART adapter at the 5’ end of the read was trimmed using TrimStartingSequence and polyA tails were trimmed using PolyATrimmer. Next, the trimmed and filtered BAM files were converted to fastq files and were used for subsequent alignment. These data can be accessed through the GEO repository (GSE161975). Reads were aligned using STAR (v.2.6.0) (30) and Ensembl (v.91) (29) annotation and the BAM file was sorted by query name using SortSam (Picard). The sorted alignment files and the unaligned (tagged) BAM files were then merged to recover BAM tags, lost during alignment (MergeBamAlignment from Picard). TagReadWithGeneFunction provides three tags for each read (gene name, gene strand and gene function) required to create a digital expression matrix. This cell matrix contains two subpopulations of cells, one cell population with many genes and reads and one with few genes and reads per cell. As the cell population with few genes and reads does not recapitulates biological signal, these needed to be removed. The average number of genes per cell (5045) clearly separated the two subpopulations, therefore, only cells with more than 5045 genes were retained (MIN_NUM_GENES_PER_CELL=5045). Furthermore, only genes with at least 2 read counts were retained. The matrices for 1/1000 and 1/10,000 diluted ERCC spikes were merged.

### Data analysis of the Chromium RNA sequencing data

Demultiplexing of the raw sequencing data was done by 10x Cell Ranger (v.2.0.2) software ‘cellranger mkfastq’ which wraps Illumina's bcl2fastq. The fastq files obtained after demultiplexing were used as input for ‘cellranger count’, which aligns the reads to the hg38 human reference genome using STAR (30) using Ensembl (v.91) (29) annotation and collapses to UMI counts. This was extended with mapping to ERCC spike-in RNA sequences, generating two separate matrices. Aggregation of samples to one dataset was done using ‘cellranger aggr’. The gene and ERCC count matrices were merged and only cells containing ERCC spikes were retained.

### Quality control and filtering of the single cell sequencing data

Quality assessment and further filtering were done in R (v.3.5.0) using Seurat (v.2.3.4) (32) and Scater (v.1.8.0) (33) as described by Lun et al. (34). For the C1 dataset, only genes with at least 5 counts were retained, as described previously (35). To retain a similar fraction of genes for the other two single cell devices, genes in at least 17 and 20 cells were retained for ddSeq and Chromium, respectively. The cyclone function of the scran (v.1.8.4) package was used to determine the cell cycle stage of the cells.

### Differential analysis of the single cell sequencing data using PIM and EdgeR-Zinger

For testing differential gene expression (DGE) between the nutlin-3 and vehicle treated cells, edgeR in combination with Zinger for the single cell experiments (36, 37) and probabilistic index models (PIM) (38) (biorxiv, DOI: 10.1101/718668) were used. Zinger calculates weights from zero-inflated negative binomial models, which is used by edgeR to fit a weighted generalized linear model (GLM) with negative binomial distribution. The PIM is a distribution-free regression model that models the probabilistic index (PI) as a function of the treatment factor. The PI indicates the degree of the difference in the distribution of gene expression between the treatment groups. PI ranges from 0 to 1 and genes are defined as upregulated in the nutlin-3 treated group if PI > 0.6 and downregulated if PI < 0.4.

### TP53 pathway activity score

Cells were ranked based the total count for TP53 pathway genes (39). In particular, cells were ranked according to the sum of log-CPM for 116 TP53 pathway genes (39). Ranks were then compared between the treatment and control group and significance was determined using the Wilcoxon rank sum test.

### Gene set enrichment analysis

Genes were ranked according to their log fold change in decreasing order and used as input for a preranked gene set enrichment analysis (GSEA) (40). The C2 (curated gene sets) gene sets were used to identify significantly enriched gene sets (q<0.05) in the datasets.

### Donwsampling of sequencing data

The *subSeq* R Bioconductor package (v 4.0) was used for downsampling of a given read count matrix (matrix ***Y*** with dimension *G*×*n*) to a new read count matrix (matrix ***X*** with dimension *G*×*n*) (41). The total number of reads of ***X*** is *p* times the total number of read counts of ***Y***, for 0 ≤ *p* ≤ 1. In particular, the subsampling procedure assumes that the read counts in ***X*** follow a binomial distribution with trial size of the total reads in ***Y*** and binomial probability of *p*.

The Chromium pseudobulk data was created by pooling of *k* cells from the Chromium data, with *k* chosen so that the number of pseudobulk samples is equal to the number of bulk samples in the NGP nutlin data. In particular, in each treatment group, cells were first shuffled and divided into groups each containing exactly *k* cells. The read counts of each gene in cells of a given group are summed to create pseudo-bulk gene expression levels. Afterwards, subsampling (as discussed above) is applied to the bulk dataset make sure that the pseudobulk and real bulk data have an equal number of total reads.

To compare the Chromium and C1 single-cell RNA-seq datasets, first, 83 cells from the Chromium data were randomly selected. In particular, the number of cells in each treatment are equal in both datasets. Afterwards, subsampling (as discussed above) is applied to make sure that reduced Chromium and C1 data have an equal number of total reads.

## RESULTS

### Experimental design

To compare single cell polyA[+] RNA-seq data generated with the C1 (Fluidigm), ddSeq (Bio-Rad/Illumina) and Chromium (10x Genomics), the same cellular perturbation experiment was performed on all three devices. Additionally, the same experiment was also performed in bulk for ten replicates to contrast with the single cell RNA-seq results (**Figure 1**). Since cell cycle status may be a confounder in single cell experiments, cell cycle synchronization by serum starvation of NGP neuroblastoma cells was carried out for all single cell experiments prior to treatment, resulting in an arrest in the G0/G1 phase (**Supplementary Figure 1A**). Next, NGP cells were treated with nutlin-3 or vehicle (ethanol). Nutlin-3 is a TP53 activator by inhibiting the interaction between TP53 and its negative regulator MDM2, resulting in an activation of the TP53 pathway and consequently in cell cycle arrest and apoptosis (1). The effect of the nutlin-3 treatment was confirmed using RT-qPCR on bulk cells and indicated a 28-fold upregulation of *CDKN1A*, a known TP53 target gene (**Supplementary Figure 1B**). ERCC spike-in RNA was added in all single cell experiments.

**Figure 1:**
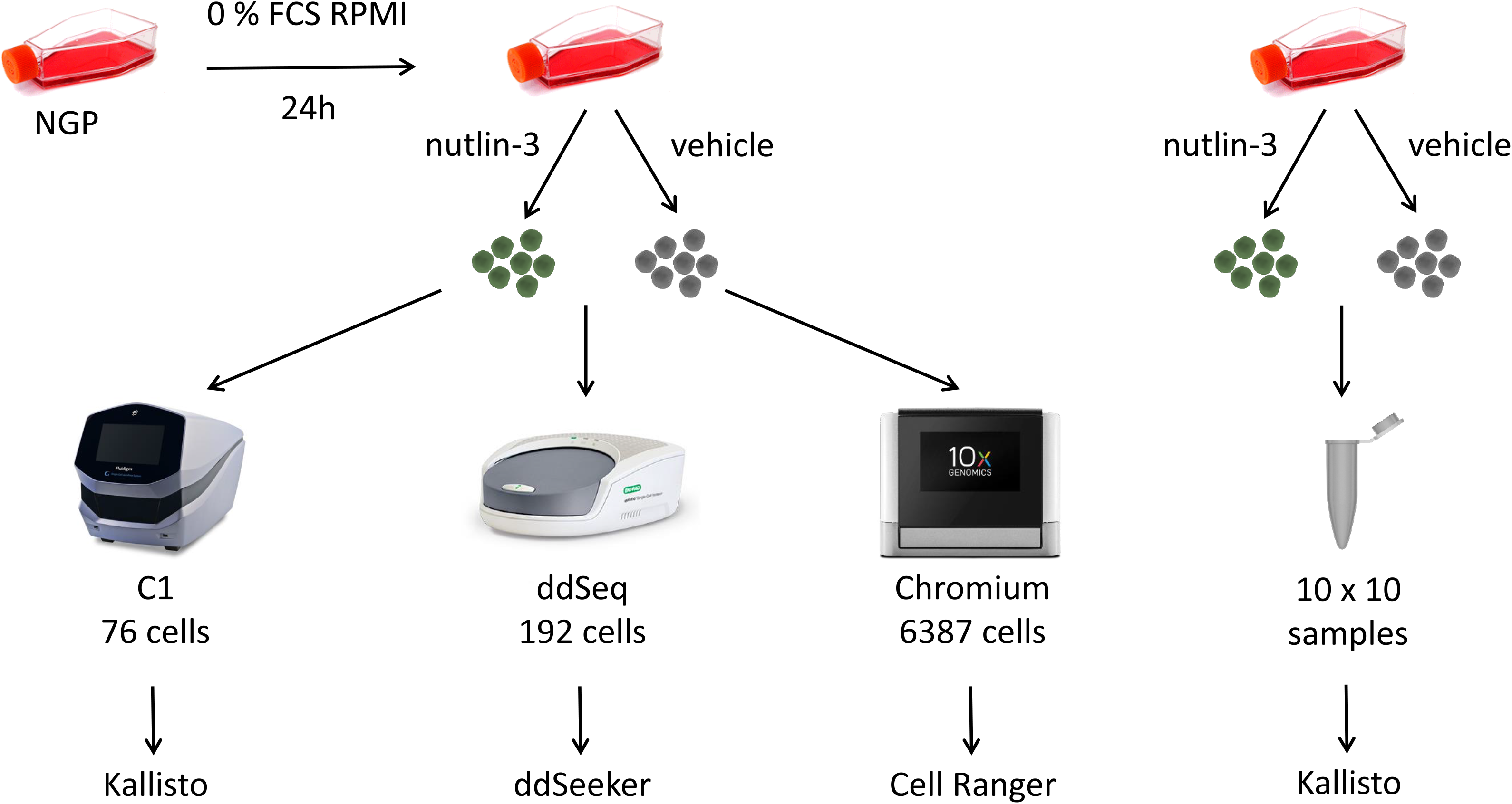
overview of the experimental set-up. Synchronized NGP cells were treated with either nutlin-3 or vehicle and single cell RNA-seq was performed using the C1, ddSeq and Chromium device. In parallel, bulk RNA-seq of 10 replicates of NGP cells treated with nutlin-3 and vehicle was carried out. Each dataset was analyzed with the appropriate pipeline.

### Quality control and filtering of sequencing data

All three single cell methods generated high quality libraries as confirmed by Bioanalyzer or Fragment Analyzer (**Supplementary Figure 1C**). Single cell RNA-seq data differ amongst others in the generated read structure, as ddSeq and Chromium reads for instance contain UMIs, while this is not the case for C1 reads. Therefore, each device has its own pipeline to analyze the data. Nevertheless, all reads, including those generated with the bulk RNA-seq protocol, were mapped against Ensembl v91, making the data comparable (**Figure 1**). For C1, the number of single cells was determined visually and 83 of the 96 capture sites contained single cells without visible debris. In contrast, single cells isolated with ddSeq and Chromium cannot be visualized and the number of single cells is determined by the computational pipeline, resulting in 260 and 7514 single cells for ddSeq and Chromium, respectively. No ERCC spikes were detected in 7 out of the 7514 Chromium isolated cells and these cells were removed from further analysis. To filter out low quality cell data, all cells with a log-transformed number of reads or genes more than three times the median absolute deviation (MAD) below the log-transformed median were removed from further analysis, since transcripts are likely not efficiently captured in these cells (2). Similarly, cells above this cut-off were also removed, as these data may be derived from cell doublets. Since we added ERCC spike-in molecules in all three single cell experiments, the same MAD cutoff was used to remove low quality cells and cell doublets based on the percentage of ERCC spike-in reads per cell. Finally, 76, 192 and 6387 single cells were retained for the C1, ddSeq and Chromium, respectively (**Table 1**, **Figure 1**). Besides low-quality cells, also genes that are only expressed in a few cells were removed. Due to the differences in throughput, the selected cut-off differs depending on the device and ~58 % of the genes were maintained by retaining only genes expressed in at least 5 (16,921 genes), 17 (12,753 genes) and 20 (15,307 genes) cells for the C1, ddSeq and Chromium, respectively. For the bulk experiment, genes expressed in fewer than three samples were removed, retaining 33,700 genes. In general, the average gene expression correlation among the platforms was high. As expected, the correlation between ddSeq and Chromium was slightly higher (r=0.84), compared to each of these methods with the C1 (ddSeq: r=0.77, Chromium: r=0.78) as ddSeq and Chromium generate sequencing libraries in a similar way (Supplementary Figure 2A). Furthermore, the average gene expression over all cells in the C1 dataset correlates best with bulk (r=0.83) (Supplementary Figure 2B), with both methods sequencing full transcripts.

**Table 1:** Overview of the number of cells removed based on library size, number of genes and percentage ERCC spikes per cell.

|  | library size | number of genes | percentage ERCC spikes | remaining cells |
| --- | --- | --- | --- | --- |
| C1 | 0 | 0 | 7 | 76 |
| ddSeq | 28 | 36 | 6 | 192 |
| Chromium | 360 | 211 | 822 | 6387 |

### The C1 has the highest gene detection sensitivity

After filtering, an average of 0.71 million, 3780 and 9466 reads were retained per cell, resulting in the detection of on average 7621, 1487 and 2220 genes per cell for the C1, ddSeq and Chromium, respectively, demonstrating that the C1 has the highest sensitivity (**Figure 2A-B**). Of note, 0.1 %, 1.5 % and 16.8 % of the reads were respectively attributed to ERCC spikes. Single cell RNA-seq experiments suffer from a lot of missing data points (dropouts) that can be biological or technical. For C1, 54.96 % of the values are dropouts, while this is much higher for ddSeq (88.34 %) and Chromium (85.50 %). PCA plots show a separation between nutlin-3 and vehicle treated cells for all single cell devices. While the distinction is clear for ddSeq, there is more overlap between treated and untreated cells for the C1 and Chromium (**Supplementary Figure 3A**). In general, genes that are low abundant are detected in a few cells, while more abundant genes are expressed in a higher fraction of cells (**Figure 2C**). ddSeq and Chromium display a tighter curve compared to the C1, probably due to the higher number of cells and removal of amplification bias by UMIs. Furthermore, ddSeq and Chromium data contain more genes that are expressed in only a few cells compared to C1, where genes are generally detected in a larger fraction of cells (**Figure 2C**). While most of the genes detected using single cell RNA-seq are also detected with bulk RNA-seq, a tiny fraction of genes is only detected by one of the devices (**Figure 3A**). In general, genes detected by all platforms display a higher expression level compared to genes detected by only one device (**Figure 3 B-D**). 16.3 % and 7.2 % of all reads map on the top 25 expressed genes for the C1 and ddSeq, respectively, while this number is higher for Chromium (29.6 %), highlighting the lower library complexity of Chromium libraries. The top 25 abundant genes contain many ribosomal and mitochondrial genes (**Supplementary Figure 3B**). Overlap analysis shows that the top 25 genes in terms of abundance differ per platform (**Supplementary Figure 3C**).

**Figure 2:**
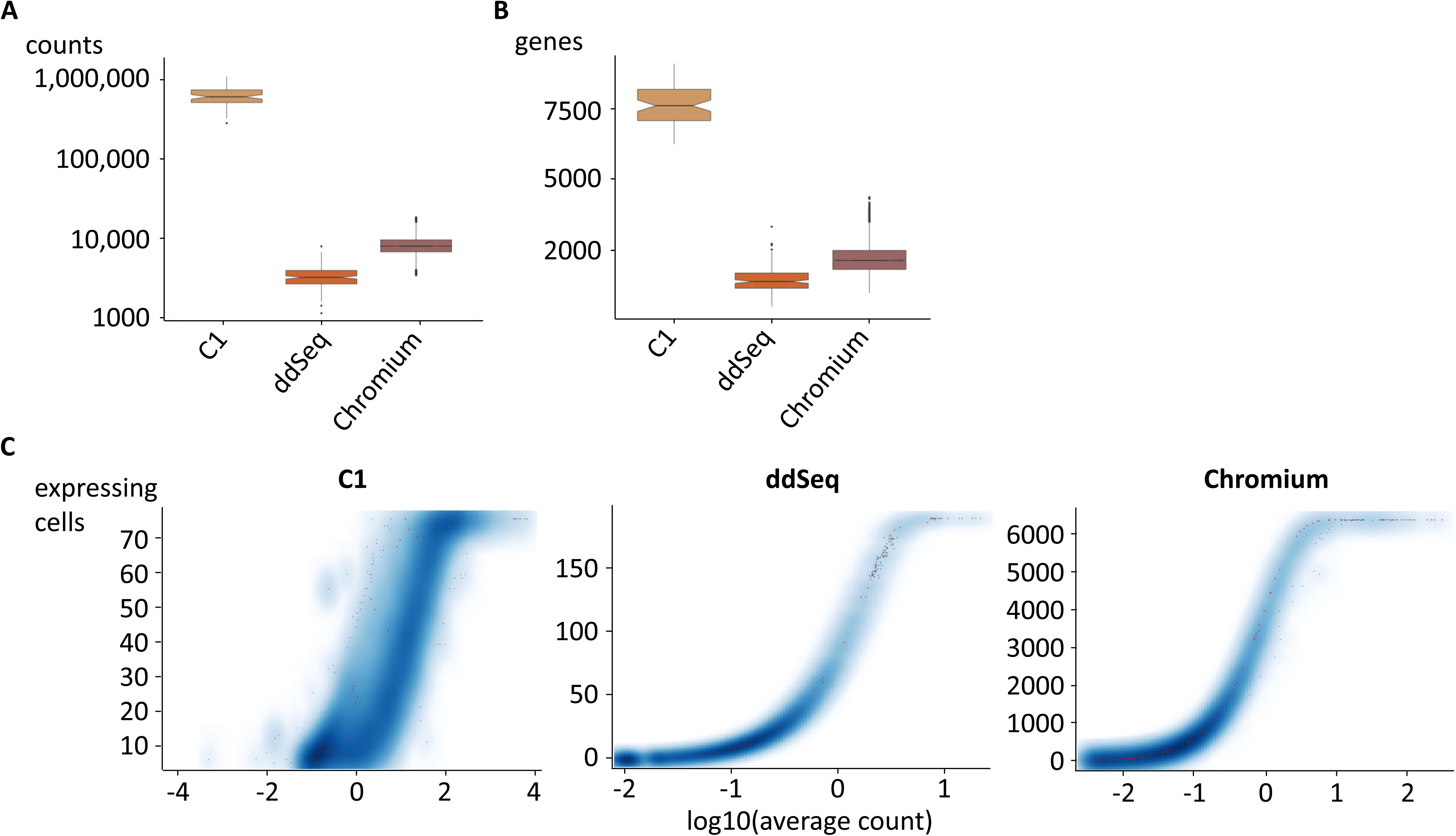
number of counts and genes detected per device. Boxplots depicting the number of counts (A) and genes (B) detected for the C1, ddSeq and Chromium after filtering. (C) Smoothscatter plot shows the correlation between the gene expression level and the number of cells that express the gene. Red dots show the ERCC spikes.

**Figure 3:**
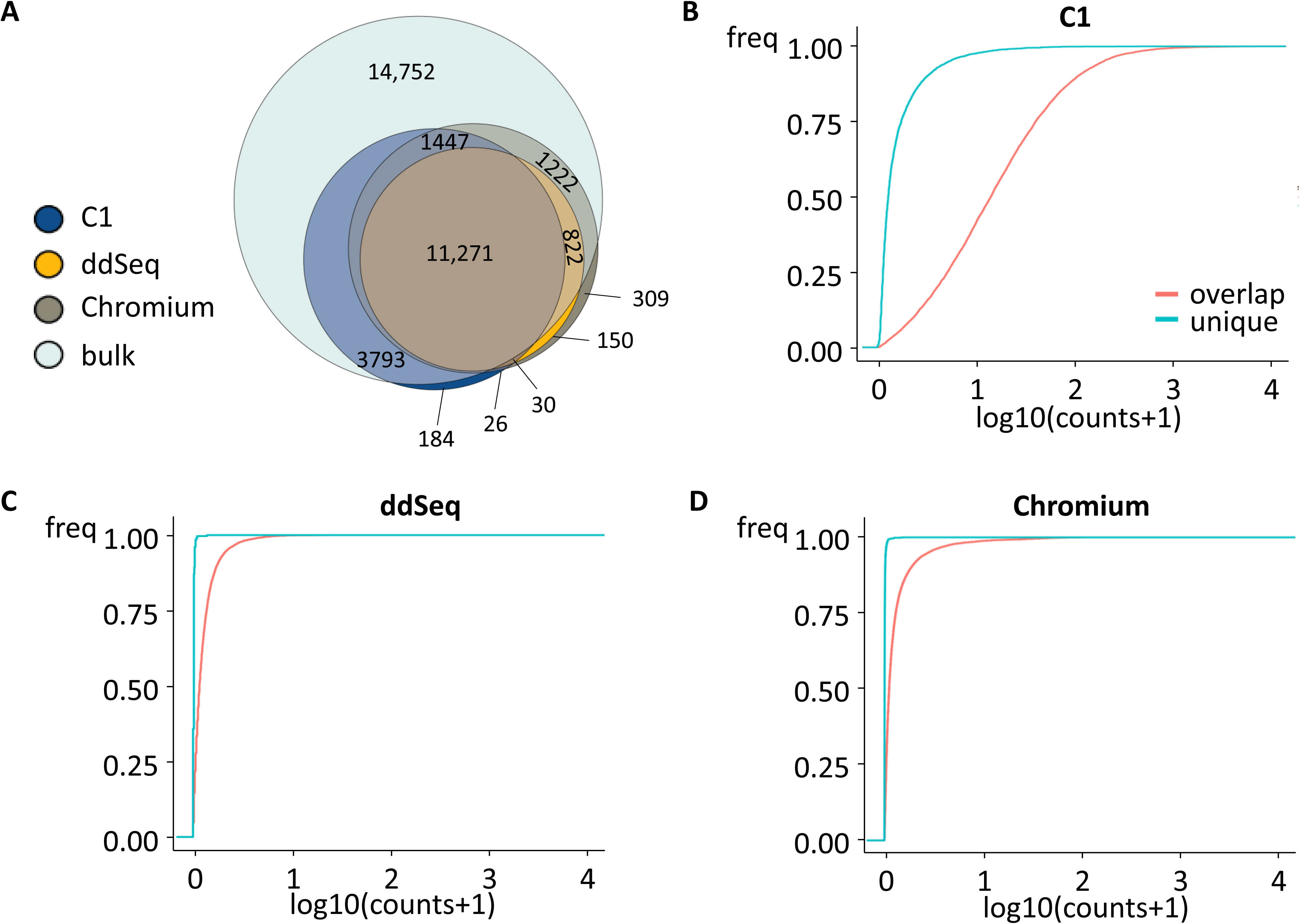
each platform detects a unique set of genes. (A) Overlap between detected genes using bulk RNA-seq, C1, ddSeq and Chromium. Cumulative expression plots of genes detected with all single cell devices or with only C1 (B), ddSeq (C) or Chromium (D).

### Bulk RNA-seq detects the largest number of differentially expressed genes, while Chromium results in most numerous enriched gene sets

As the number of differentially expressed genes in part depends on the statistical tool, we performed both EdgeR in combination with Zinger as well as probabilistic index model (PIM) analysis and retained high-confident consensus genes that are called significantly differentially expressed with both tools (3–5). In a comparison study, EdgeR was shown to be one of the better tools for single cell differential gene expression analysis and PIM is a new tool, developed specifically for differential gene expression analysis of single cells (biorxiv, DOI: 10.1101/718668). Genes were called significantly differentially expressed by EdgeR if FDR < 0.05 and absolute log fold change > 1, while genes are significantly differentially expressed according to PIM if adjusted p-value < 0.05 and PI < 0.4 (downregulated) or PI > 0.6 (upregulated). For bulk, C1, ddSeq and Chromium, 7010, 40, 28 and 88 significantly differentially expressed genes were identified, respectively (**Supplementary Table 2**). By only including genes that are detected by all four platforms, the number of differentially expressed genes in the bulk dataset drastically dropped to 1665, while only little differences were noticed for C1 (36 genes), ddSeq (28 genes) and Chromium (86 genes), in line with the fact that many genes are only detected in the bulk experiment. While most differentially expressed genes in the single cell datasets overlap with those detected in the bulk dataset, some genes are uniquely differentially expressed in only one of the datasets (**Figure 4A**). Genes that are differentially expressed in only one of the single cell datasets are mostly borderline in significance and effect size (**Supplementary Figure 4**). Interestingly, although many more genes are significantly differentially expressed in the bulk dataset compared to the single cell datasets, enrichment analysis shows that Chromium identifies more significantly (q-value <0.05) positively enriched gene sets, demonstrating that biological signal can be effectively captured with a subset of the most abundant genes (**Supplementary Table 2**, **Figure 4B**). Of note, several TP53 gene sets pop up in all positively enriched gene sets, and cell cycle gene sets are common in the negatively enriched gene sets, validating the effect of nutlin-3 on the TP53 pathway and the cell cycle arrest in nutlin-3 treated cells for all datasets. Furthermore, the TP53 activity scores are significantly different (p-value < 0.01) in nutlin-3 treated cells compared to vehicle treated cells for all devices, supporting the fact that all methods are capable of detecting biological signal (**Figure 4C**). Of note, the bulk experiment has the clearest separation between treated and untreated cells, but this may be in part due to the fact that TP53 target genes were previously defined based on bulk gene expression profiles (6).

**Figure 4:**
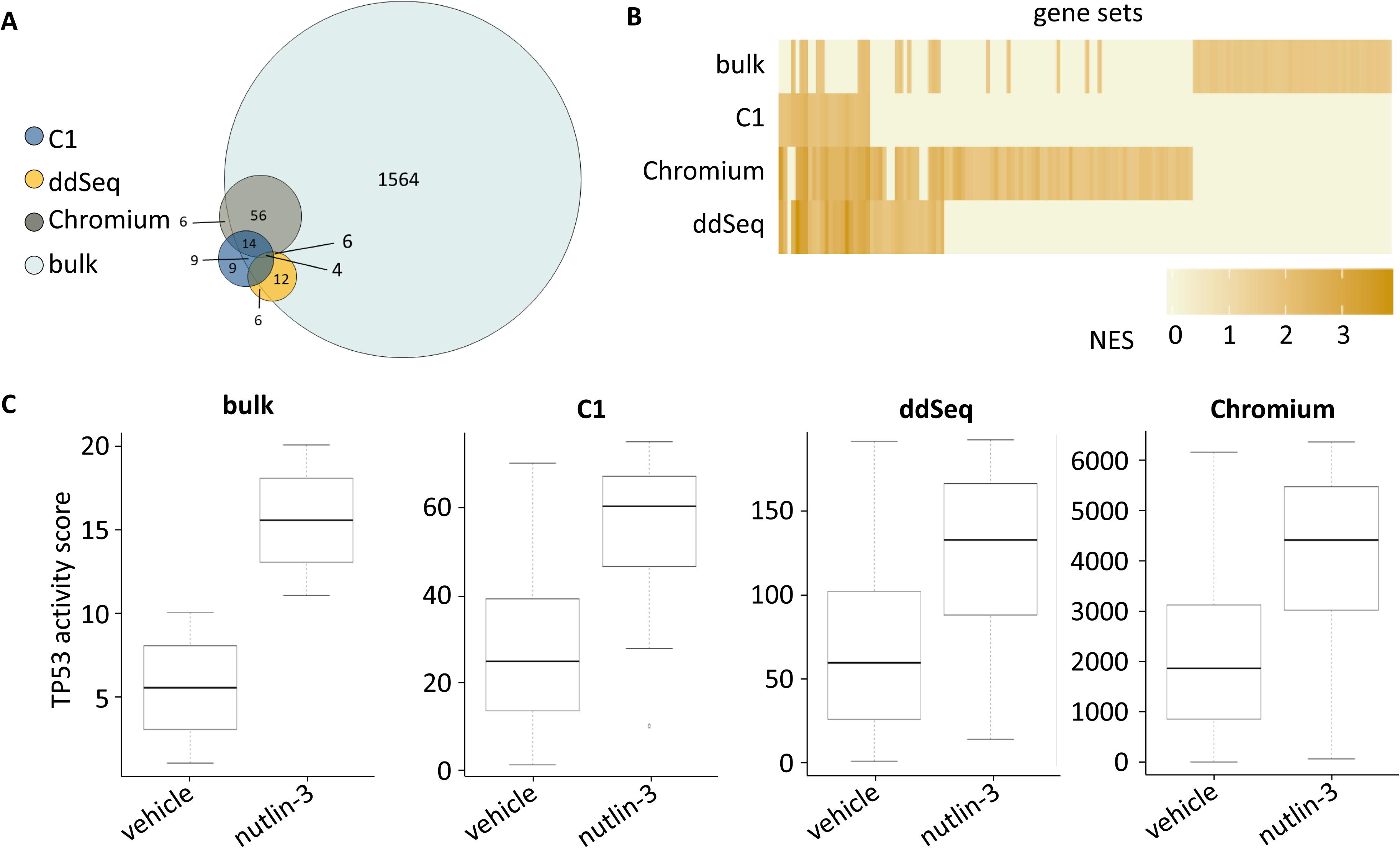
single cell RNA-seq results recapitulate the biological signal. (A) Overlap of genes detected by all platforms and significantly differentially expressed with both EdgeR in combination with Zinger as well as PIM. (B) Heatmap of significantly positively (q-value <0.05) enriched gene sets after GSEA for the C2-curated gene sets for each method. Gene sets are color-coded according to their normalized enrichment score (NES). (C) Boxplots depicting the TP53 activity score per cell, whereby ranking was based on the expression of 116 TP53 target genes.

### Single cell RNA sequencing reveals a heterogeneous response upon nutlin-3 treatment and uncovers hidden biological signals

To get a first view on the transcriptional response of NGP cells on nutlin-3 treatment, the expression of *CDKN1A,* a known TP53 target, was determined for the three single cell and the bulk RNA-seq experiments. While *CDKN1A* is significantly upregulated in all datasets upon nutlin-3 treatment, there is a remarkable heterogeneity of *CDKN1A* expression in the single cell datasets (**Figure 5A**). To understand the differences between cells with a low and high expression of *CDKN1A*, nutlin-3 treated cells with *CDKN1A* expression in the lowest quartile were compared to cells with expression in the highest quartile. To have a sufficiently large number of cells in each group, this analysis was only done for the Chromium dataset. In total, 83 genes were significantly differentially expressed, of which 76 overlapped with the set of genes significantly differentially expressed between nutlin-3 and vehicle treated cells in the full Chromium dataset (**Supplementary Table 2**). In addition, 93 of the 103 significantly positively enriched gene sets overlap with those of the full Chromium dataset (**Figure 5B-C**, **Supplementary Table 2**). These results demonstrate that the same signals can be detected between vehicle and nutlin-3 treated cells and between nutlin-3 treated cells with low and high *CDKN1A* expression. To validate these results, this analysis was also performed for *PUMA*. *PUMA* is another TP53 target gene that is significantly upregulated in all datasets upon nutlin-3 treatment and for which there is a remarkable heterogeneity of expression in the single cell datasets (**Supplementary Figure 5A**). 86 genes were significantly differentially expressed between the cells with an expression of *PUMA* in the lowest and highest quartile (**Supplementary Table 2).**Of these, 77 overlapped with the full Chromium dataset. 79 of the 100 significantly positively enriched gene sets overlap with those of the full Chromium dataset (**Supplementary Figure 5B-C**, **Supplementary Table 2**). These results confirm that the same signals can be detected between vehicle and nutlin-3 treated cells and between nutlin-3 treated cells with low and high expression of a TP53 target gene.

**Figure 5:**
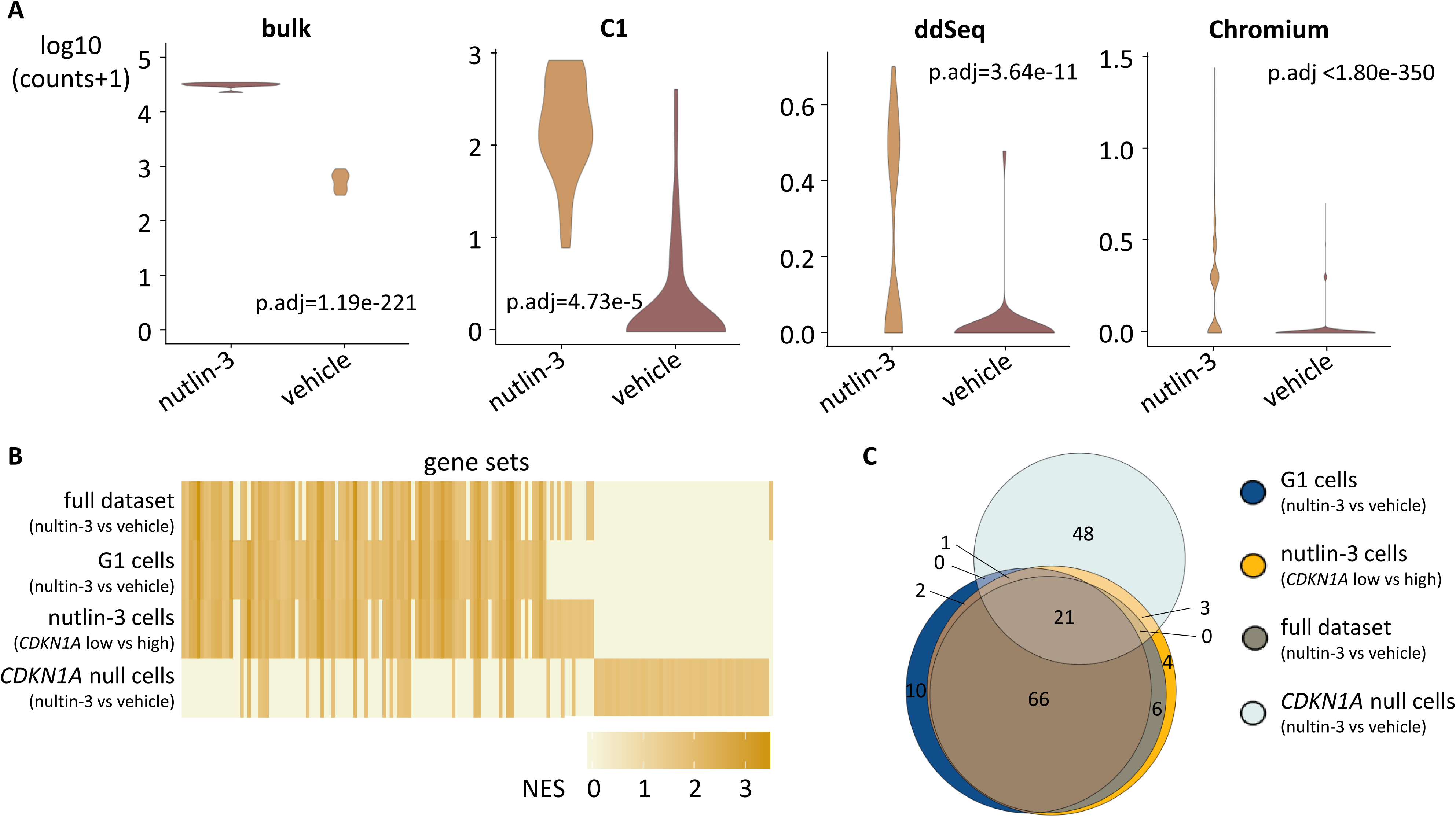
differences between cells with varying *CDKN1A* expression. (A) *CDKN1A* expression in the bulk and single cell RNA-seq datasets. (B) Heatmap of significantly positively (q-value <0.05) enriched gene sets after GSEA for the C2-curated gene sets for each subgroup. Gene sets are color-coded according to their normalized enrichment score (NES). (C) Overlap between significantly positively enriched gene sets for the three cellular subgroups and full Chromium dataset.

As the cell cycle status can be a major confounder in single cell experiments masking putative biological effects, cells in the G1 phase of the cell cycle in the Chromium data set were selected based on the expression of a G1 cell cycle signature. Doing so, 105 genes were significantly differentially expressed between nutlin-3 and vehicle treatment, of which 80 overlapped with the differentially expressed genes of the full Chromium dataset (**Supplementary Table 2**). As we detected slightly more (105 instead of 88) differentially expressed genes between nutlin-3 and vehicle treated cells in the G1 phase compared to the full dataset, these differentially expressed genes might have been masked by cell cycle effects in the full dataset. Several genes that are downregulated in the G1 cells, but not in the full dataset, including *UBE2C* and *PCLAF,* are known to be repressed by TP53 and also downregulated in the bulk RNA-seq dataset (7, 8). Likewise, several genes that are upregulated in the G1 cells only, including *DDIT4* and *KRT17,* are known to be induced by TP53 and also upregulated in the bulk RNA-seq dataset, showing that more biologically relevant targets are identified in RNA-seq data from single cells in the same cell cycle phase (9, 10). Interestingly, *PTTG1* is significantly differentially expressed in G1 cells, but not in the full Chromium, nor the bulk dataset, and known to be repressed by TP53 (11). Additionally, 87 of the 100 significantly positively enriched gene sets overlap with those of the full dataset (**Figure 5B-C**). One gene set (CONCANNON_APOPTOSIS_BY_EPOXOMICIN_UP, NES= 1.80, FDR = 0.03) containing genes upregulated because of apoptosis was only enriched in the G1 cells, supporting the relevance of signals that are only detected in the G1 cells and not in the full dataset.

Finally, single cell experiments are characterized by a high dropout rate. To determine the differences between nutlin-3 and vehicle treated cells without detectable expression of *CDKN1A,* such so-called *CDKN1A* null cells were selected for each treatment arm. 71 genes were significantly differentially expressed, of which 68 overlapped with the full set (**Figure 5B-C**, **Supplementary Table 2**). In contrast, only 21 of the 73 significantly positively enriched gene sets overlap with those of the full dataset. Validating these results for *PUMA* null cells revealed similar to *CDKN1A* null cells a large overlap in the significantly differentially expressed genes (70 of the 72 genes). In contrast to *CDKN1A* null cells, there was a large overlap of the significantly positively enriched gene sets (71 of the 81) with the full dataset, highlighting the complexity of dropouts in single cell experiments.

### Pseudobulk data resembles real bulk data better than single cell data

To understand the differences between bulk RNA-seq and single cell RNA-seq patterns better, pseudobulk data from the single cell data were created by pooling and averaging subsets of single cells. Chromium data were pooled in ten pseudosamples per treatment arm, resulting in the same sample size as the bulk data. Chromium was taken as an example as this dataset contains the highest number of cells. To make the data even more comparable, the bulk library size was downsampled to obtain the same number of reads for the single cell and bulk dataset summarized over all (pseudo)samples. Originally, the bulk library size was 4.8 times larger compared to the single cell library size. After downsampling, the total number of reads in each experiment was 70.2 million, with a mean of 3.5 million reads per (pseudo)sample (**Figure 6A**). As done before for the original bulk data, only genes expressed in at least 3 samples were retained in the downsampled bulk and pseudobulk datasets. The correlation between the average gene expression in the downsampled bulk and pseudobulk dataset was higher (r=0.83) compared to the correlation in the original bulk and single cell dataset (r=0.69) (**Supplementary Figures 2 and 6A**). As expected, fewer genes (27,659 instead of 33,700) and fewer significantly differentially expressed genes (4606 instead of 7010) were detected in the downsampled bulk dataset compared to the original bulk dataset, due to the lower sequencing depth (Supplementary Table 2). Of these differentially expressed genes, the large majority (4012, 87.10 %) overlapped with the differentially expressed genes of the original bulk dataset. For the pseudobulk dataset, almost 10-fold more genes (810 instead of 88) were significantly differentially expressed compared to the original single cell dataset. Of note, this number is still considerably lower compared to bulk at equal sequencing depth. The higher number of differentially expressed genes in the pseudobulk dataset compared to the original single cell dataset is probably owing to the reduction in noise after pooling single cells into pseudobulk samples. 509 of the 810 significantly differentially expressed genes in the pseudobulk dataset overlap with the differentially expressed genes of the downsampled bulk dataset. Furthermore, 123 and 101 significantly positively enriched datasets were identified for the pseudobulk and downsampled bulk dataset, respectively. With 42 of the 123 positively enriched gene sets in the pseudobulk dataset overlapping with those of the downsampled bulk, and only 18 of the 94 positively enriched gene sets overlapping in the original single cell data and the bulk data, the pseudobulk data better resembles the bulk data (**Figure 6B**, **Supplementary Figure 6B**). Furthermore, the TP53 activity scores are significantly different (p-value < 0.01) in nutlin-3 treated cells compared to vehicle treated cells for both the downsampled bulk and pseudobulk dataset, showing that pseudobulk samples continue to recapitulate biological signal. Interestingly, there is a clearer separation for the pseudobulk dataset compared to the original single cell dataset (Figure 4C, 6C).

**Figure 6:**
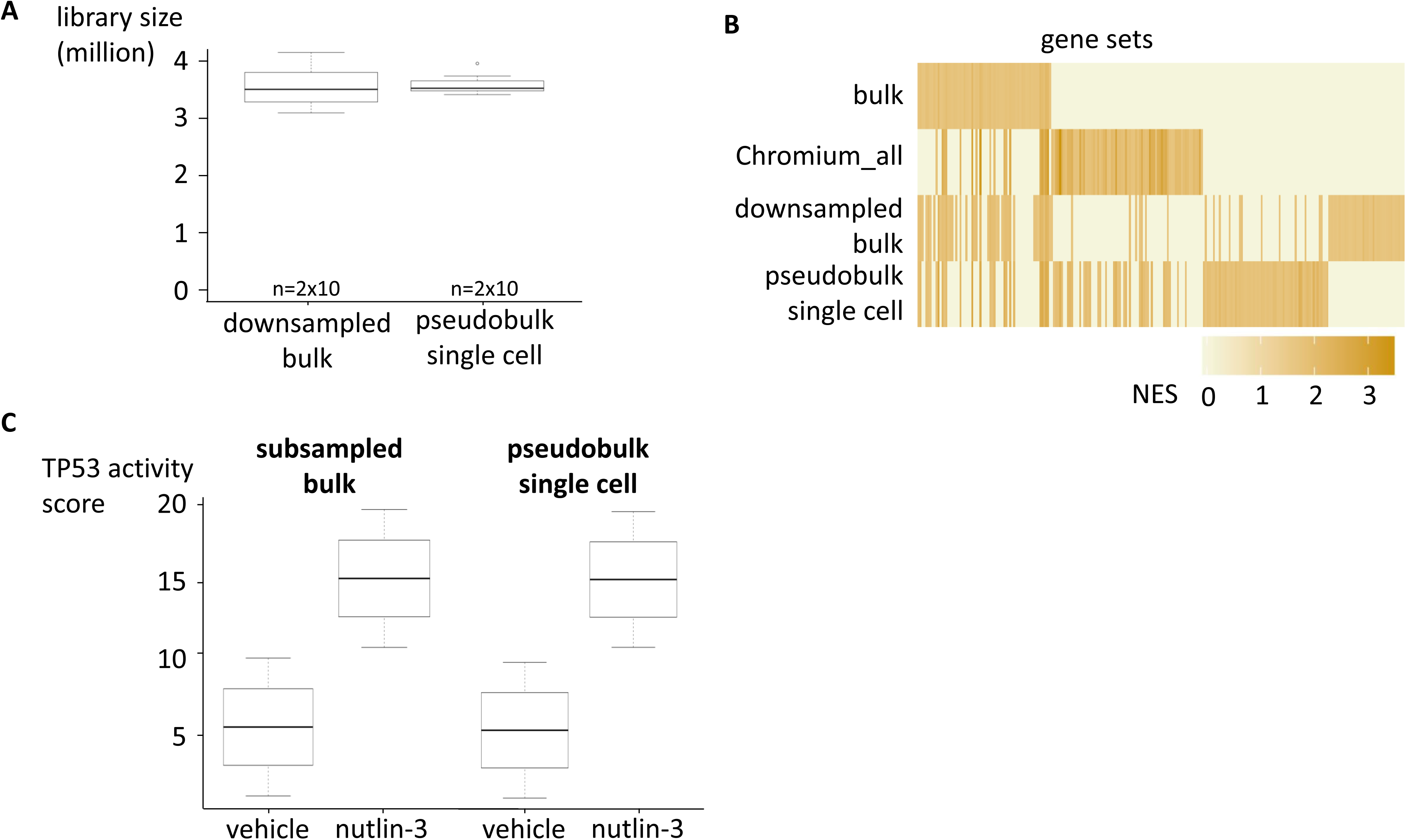
downsampling of bulk and pseudobulkification of single cell data. (A) After downsampling of the bulk data and generating pseudobulk data from the Chromium single cell RNA-seq data, the mean number of reads per (pseudo)sample is 3.5 million. (B) Heatmap of significantly positively (q-value <0.05) enriched gene sets after GSEA for the C2 curated gene sets for the original and downsampled bulk and pseudobulk Chromium datasets. (C) Boxplots depicting the TP53 activity score per cell, whereby ranking was based on the expression of 116 TP53 target genes.

### The C1 and Chromium datasets become more comparable when comparing an equal number of cells and equal sequencing depth

To determine the effect of the sequencing depth on single cell experiments, 83 cells of the Chromium dataset were randomly selected to compare with the 83 cells of the C1 dataset. To make the datasets even better comparable, the C1 dataset was downsampled to the sequencing depth of the subset of 83 cells of the Chromium dataset. Originally, the Chromium library size of the complete dataset was 1.17 times larger compared to the C1 dataset. After selecting an equal number of cells for the Chromium and C1 dataset and downsampling the C1 dataset, the total number of reads in each experiment was 0.8 million, with a mean of 9652 and 9645 reads per sample for Chromium and C1, respectively (**Supplementary Figure 7A**). Only genes expressed in at least 3 samples were retained in both datasets. The correlation between the average gene expression in the downsampled C1 dataset and subset of the Chromium dataset was somewhat lower (r=0.73) compared to the correlation in the original C1 and Chromium single cell datasets (r=0.78) (**Supplementary Figures 2 and 6C**). Surprisingly, more significantly differentially expressed genes (71 instead of 40) were detected in the downsampled C1 dataset compared to the original C1 dataset (**Supplementary Table 2**). However, of these differentially expressed genes, only 21 overlapped with the differentially expressed genes of the original C1 dataset. For the subset of the Chromium dataset, fewer genes (79 instead of 88) were significantly differentially expressed compared to the original single cell dataset, probably due to the reduction in cell number. Only 10 of the 71 significantly differentially expressed genes in the subsampled C1 dataset overlap with the differentially expressed genes of the subset of the Chromium dataset. Furthermore, 21 and 53 significantly positively enriched datasets were identified in the subsampled C1 and subset of Chromium dataset, respectively. With 18 of the 53 positively enriched gene sets in the subset of the Chromium dataset overlapping with those of the downsampled C1 dataset (Jaccard 0.39), and 20 of the 94 positively enriched gene sets of the original Chromium dataset overlapping with the original C1 data (Jaccard 0.21), it is clear that by taking an equal number of cells for both datasets and subsampling the data, the Chromium and C1 dataset are more similar (**Supplementary Figure 6D-7B**). Furthermore, the TP53 activity scores remain significantly different (p-value < 0.01) in nutlin-3 treated cells compared to vehicle treated cells for both the subsampled C1 and subset of Chromium dataset, indicating that these smaller data-sets continue to recapitulate biological signal (**Supplementary Figure 7C**).

## Discussion

Over the last years, several single cell RNA-seq methods emerged, whereby the number of single cells drastically increased from a few dozen up to tens of thousands of single cells. While several studies attempted to compare these single cell RNA-seq methods, most studies focused on the quality of the generated data and their ability to distinguish cellular subpopulations (1–5). Furthermore, the more recent ddSeq instrument was included in only one comparative study (2). Here, we evaluated for the first time three commercially available single cell devices - C1, ddSeq and Chromium - to study transcriptional heterogeneity upon a therapeutic perturbation and to contrast it with a bulk cell population response. To this purpose, NGP neuroblastoma cells were treated with the TP53 activator nutlin-3 or vehicle as negative control followed by single cell RNA-seq using the C1, ddSeq and Chromium. Since the cell cycle state is a known confounder in single cell experiments, this effect was minimized by synchronizing cells prior to treatment. To further characterize the results of the single cell experiments, bulk RNA-seq was performed in parallel on the same model system in ten biological replicates. We showed that the highest number of genes with lowest dropout rates are detected using the C1, confirming that this platform has the highest detection sensitivity, which may in part result from the higher sequencing depth (5). While downsampling read depth to an equal average number of reads per cell for all three devices should be carried out to effectively confirm that the C1 displays the highest sensitivity, different read distribution along the transcript (i.e. entire transcript (C1) vs. 3’ end only (ddSeq, Chromium)) may continue to confound such an analysis. We also observed that the overlap between the detected genes in the bulk and single cell datasets was highest for the C1 with an overlap of more than 50 %, which is slightly higher than reported previously (5). The C1 average gene expression levels correlated better with bulk gene expression data compared to ddSeq and Chromium, owing to the higher sequencing depth and higher transcriptome complexity of the C1 cDNA libraries. In contrast, correlation of average gene expression among the single cell devices revealed a slightly better correlation between the ddSeq and Chromium, in line with their similarity in terms of RNA-seq library preparation. The correlation between C1 and Chromium was the lowest, as previously reported (5, 6). The gene expression profiles of the ddSeq and Chromium seem less noisy compared to the C1, owing to the higher number of isolated single cells and the use of UMIs. It has been reported that technical noise can be reduced by 50 % using the UMI enabled counting of cDNA molecules (7, 8). Although a large overlap in the genes quantified with the three single cell devices was seen, each device also detected some unique genes. It has been previously reported that unique C1 genes do not have 3’ ends that are difficult to capture, preventing their detection by 3’ end sequencing technologies such as the ddSeq and Chromium. Hence, the large set of unique C1 genes results from higher C1 mRNA capture efficiency (1, 5). In our attempt to make the devices more comparable, ERCC spike-in RNA molecules were added to all three single cell experiments. Of note, ERCC spikes are generally not added to droplet-based experiments, since these spikes-in molecules are added to every droplet and consequently also amplified and sequenced in droplets without cells, increasing the sequencing costs considerably (1). Due to the lack of guidelines for droplet-based experiments, too many reads (17 %) in our Chromium dataset mapped to ERCC spikes, consequently losing endogenous reads and indicating that lower amounts of ERCC spike-in RNA should be added in future experiments. Apart from being used as workflow control, ERCC spike-in molecules can also be used for normalization, although this use is still under debate (9–11).

To test the ability to identify differentially expressed genes upon nutlin-3 treatment in single cell RNA-seq datasets, two different statistical methods were used, i.e. EdgeR in combination with Zinger, and PIM. As differential gene expression analysis tools typically vary in the number of genes called as differentially expressed, we here continued with the intersection of both tools to conservatively identify truly differentially expressed genes (12). The largest number of differentially expressed genes was detected in the Chromium dataset, in line with the observation that more genes are called differentially expressed with increasing number of single cells (5, 12). Although many more genes were differentially expressed in the bulk dataset, the biological signal is faithfully recapitulated in the tested single cell datasets as strong enrichment of several TP53 gene sets was present in all datasets. This result suggests that detecting the most abundant genes (through single cell RNA-seq data) is sufficient for pathway activity analysis. Of note, single cell datasets also reveal some unique enrichment signals, of which the relevance should be determined by further investigation.

To characterize the effect of nutlin-3 treatment at the single cell level, five cell subpopulations from the full Chromium dataset were selected based on their cell cycle stage and TP53 transcriptional target gene *CDKN1A* and *PUMA* expression levels, and compared with the entire cell population. These subpopulation analyses were only performed for the Chromium dataset in order to have a sufficient number of cells per subset. In order to avoid cell cycle effects as much as possible, nutlin-3 and vehicle treated cells in the G1 phase were selected in the first subset. Although a large fraction of the differentially expressed genes in the G1 cells overlapped with the full dataset, more significantly differentially expressed genes were detected in the G1 cells, possibly hidden by cell cycle effects in the full dataset. Many of these genes are known to be regulated by TP53, showing the utility of cell subpopulation analysis and the relevance of the genes differentially expressed in cells in the G1 phase. This type of subpopulation analysis could in principle be extended to the other cell cycles stages. In a second subset, differential gene expression and gene set enrichment analysis on nutlin-3 treated cells with low or high expression of the TP53 target genes *CDKN1A* or *PUMA* revealed that these subsets resemble vehicle and nutlin-3 treated cells from the full dataset. This indicates that treated cells with low expression of *CDKN1A* or *PUMA* are similar to vehicle treated cells and may thus represent cells that react in a later stage to nutlin-3 or show primary resistance. To further investigate this intriguing observation, time-course experiments should be set up to reveal if *CDKN1A* or *PUMA* is upregulated in a larger fraction of cells at later timepoints, in line with a delayed nutlin-3 response. In the third subset, nutlin-3 and vehicle treated cells without *CDKN1A* or *PUMA* expression were compared. With exception of the enrichment analysis in the CDKN1A nulls cells, overall our results indicate that nulls cells are very similar than the full population of cells. This strongly suggests that these cells likely not reflect a different subpopulation, but are – at least to a large degree- null cells because of technical dropout. While we cannot exclude that truly different null cells exist, it seems to caution us that the absence of a marker does not necessarily denote a different cell status or cell type. While our analyses provide first insights in the heterogeneous response of NGP cells on nutlin-3 treatment, more in depth analyses are required to better understand these observations.

Since single cell and bulk RNA-seq experiments differ at several points, such as the library prep method, the sequencing depth, and the ‘sample’ size, we set up an additional analysis in which we attempted to cancel out these differences. To account for the sample size, Chromium single celle data were pooled to create 10 pseudobulk samples for each condition. In addition, the bulk dataset was downsampled to obtain the same number of reads as the single cell RNA seq dataset. Correlation analysis between the gene expression profiles of the pseudobulk and bulk samples depicted a higher correlation compared to the correlation between the original bulk and the Chromium single cell dataset, validating that pseudobulk data resembles bulk data. More genes were differentially expressed upon pooling, likely because of a reduction of measurement noise, which is typically high in single cell experiments (7, 13). Still, the number of differentially expressed genes is lower compared to the bulk dataset, owing to the marked higher detection sensitivity of bulk RNA-seq methods. In addition, to make the C1 and Chromium single cell datasets more comparable, an equal number of cells was compared (n=83) at equal sequencing depth. Despite the lower correlation between the subsampled C1 dataset and the subset of the Chromium dataset compared the original datasets, more significantly positively enriched gene sets overlap, showing that analyzing an equal number of cells at equal sequencing depth, makes the data more similar.

In conclusion, we evaluated for the first time three commercial single cell RNA-seq devices in terms of their ability to characterize a cellular perturbation system. We revealed that despite the lower number of differentially expressed genes in single cell RNA-seq experiments compared to bulk RNA-seq experiments, the biological signal can be faithfully detected through gene set enrichment analysis for all single cell devices. We also demonstrated that single cell RNA-seq analyses reveal transcriptional heterogeneity in response to nutlin-3 treatment, which may form the basis to identify potentially late-responders or resistant cells that are hidden in bulk RNA-seq experiments.

## Notes

### Competing Interest Statement

The authors have declared no competing interest.

