## Supplementary Figures and Tables for "Comprehensive benchmarking of single cell RNA sequencing technologies for characterizing cellular perturbation"

### Supplementary Figure 1

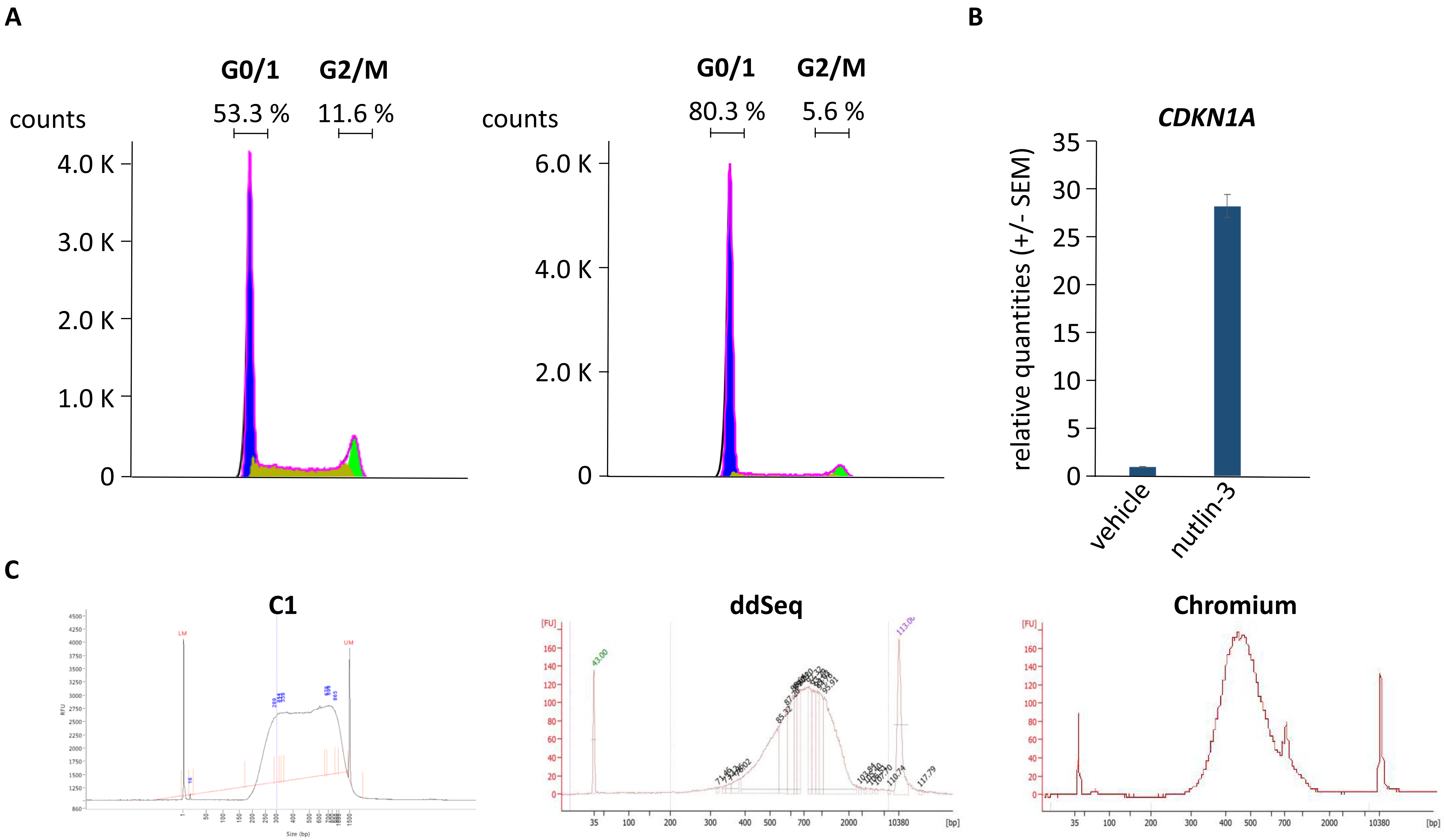

### Supplementary Figure 2

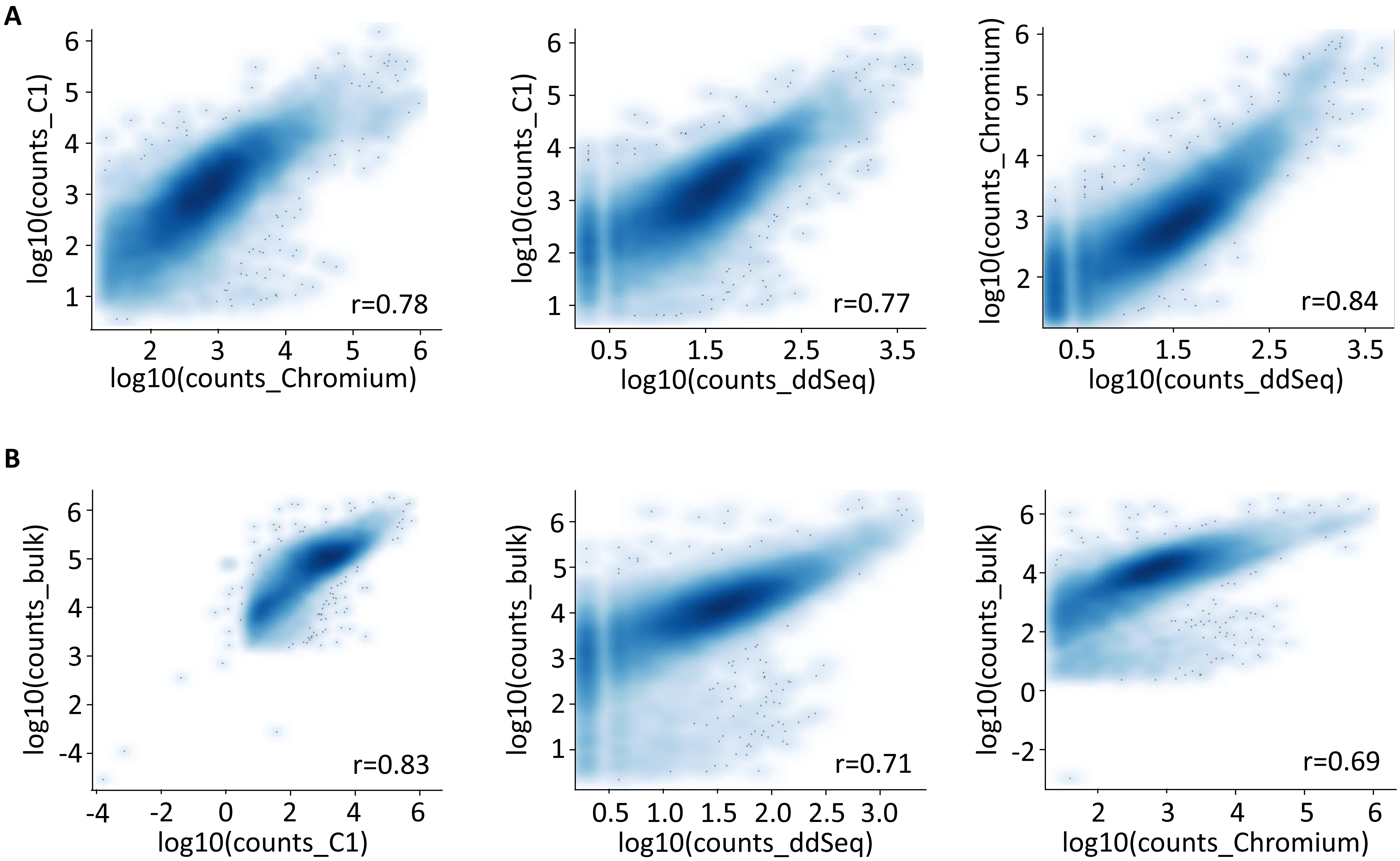

### Supplementary Figure 3

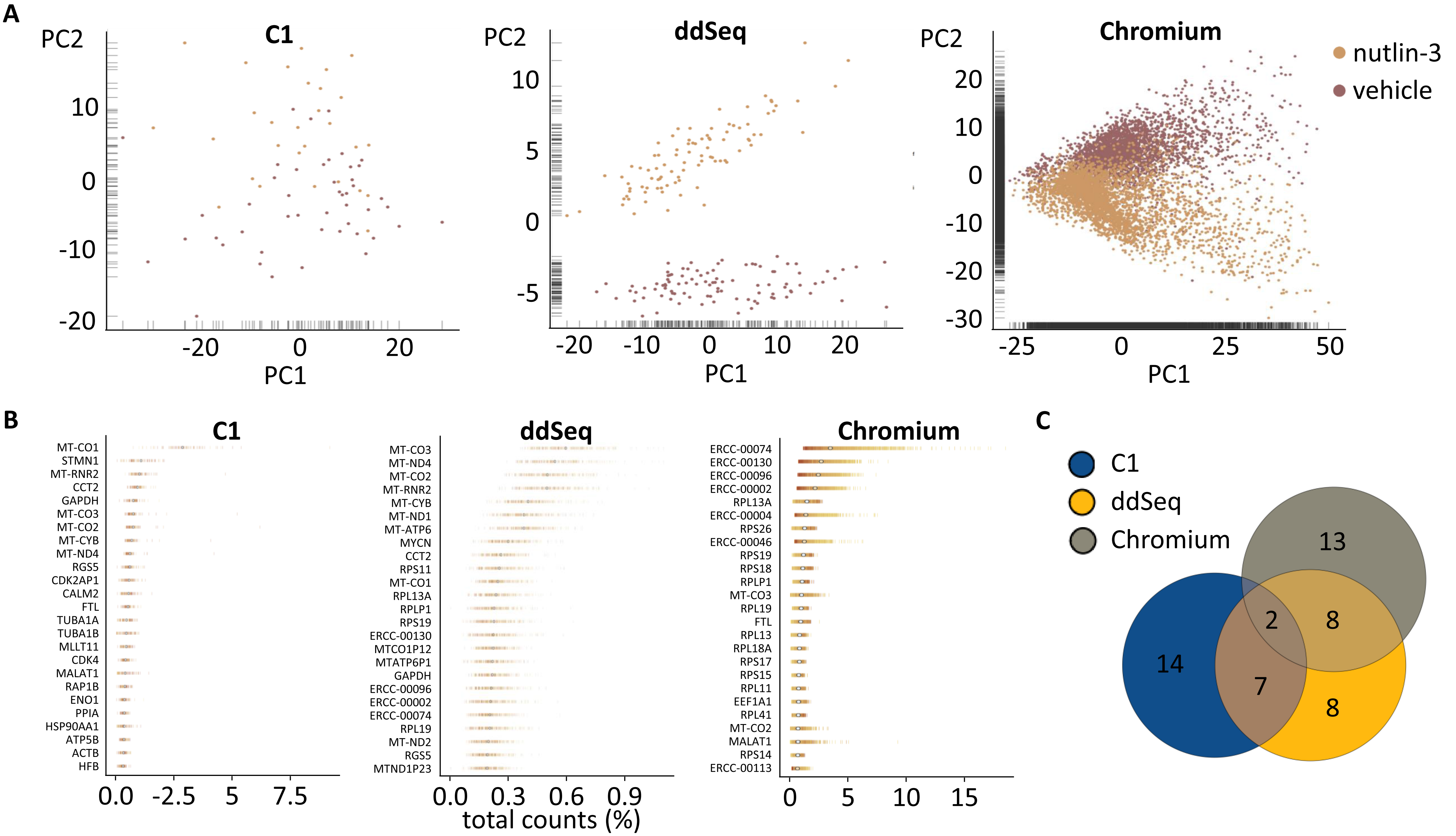

### Supplementary Figure 4

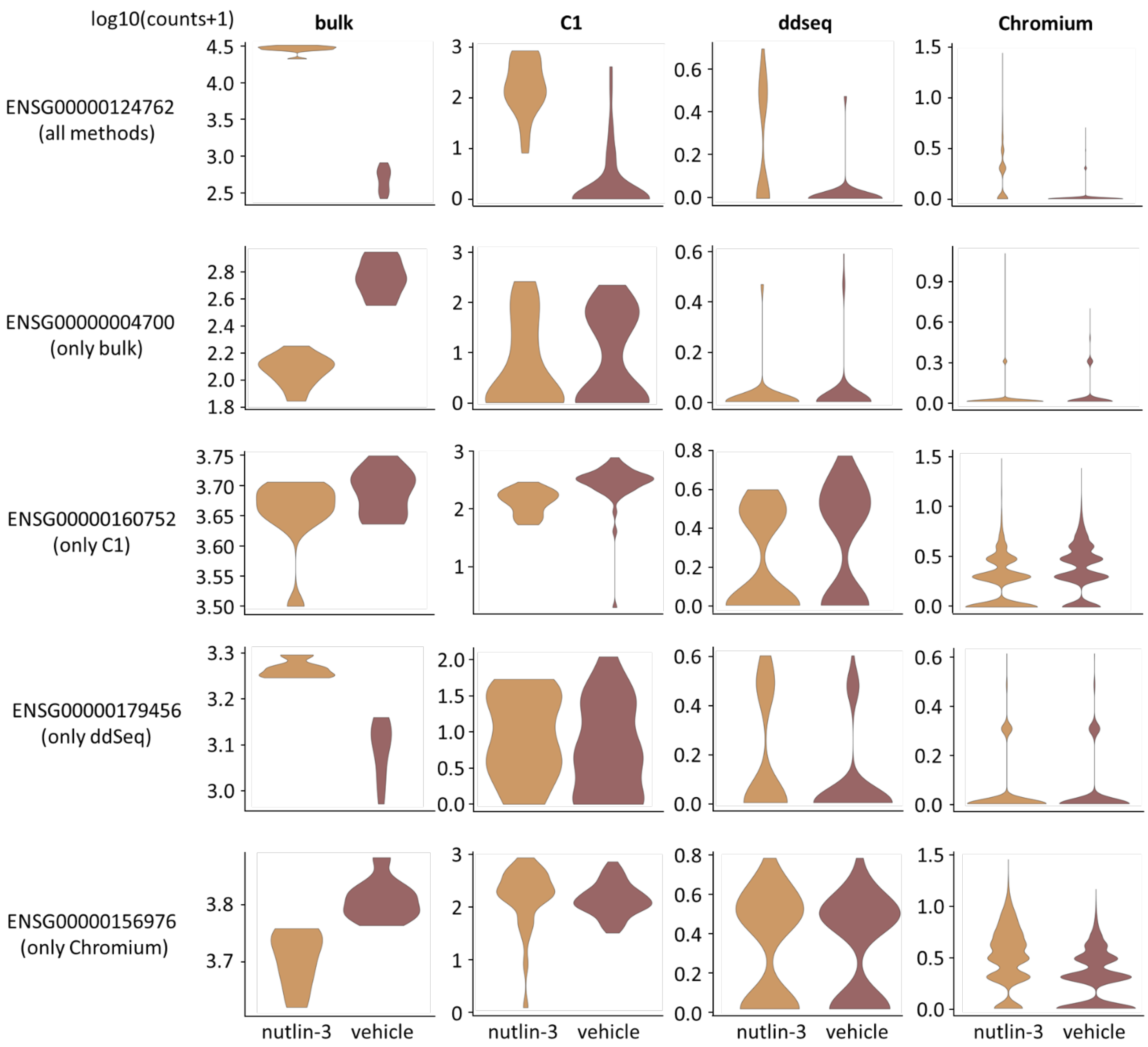

| | bulk | | | | C1 | | | | ddSeq | | | | Chromium | | | |
|---|---|---|---|---|---|---|---|---|---|---|---|---|---|---|---|---|
| | PI | p.adj | LFC | FDR | PI | p.adj | LFC | FDR | PI | p.adj | LFC | FDR | PI | p.adj | LFC | FDR |
| ENSG00000124762 | 1.00 | 1.19e-221 | 5.71 | 1.81e-321 | 0.97 | 4.73e-5 | 3.93 | 2.00e-3 | 0.73 | 6.64e-11 | 3.14 | 3.37e-18 | 0.79 | 0.00 | 2.33 | 0.00 |
| ENSG00000004700 | 1.37e-10 | 0.00 | -1.12 | 1.54e-60 | 0.43 | 0.77 | -0.10 | 0.99 | 0.49 | 0.87 | -0.42 | 0.91 | 0.45 | 5.73e-10 | -0.50 | 5.83e-15 |
| ENSG00000160752 | 0.57 | 0.78 | -0.12 | 0.07 | 0.09 | 7.93e-6 | -1.10 | 4.18e-5 | 0.42 | 0.58 | -0.42 | 0.33 | 0.40 | 1.41e-43 | -0.52 | 1.69e-43 |
| ENSG00000179456 | 1.00 | 1.94e-167 | 0.57 | 5.30e-35 | 0.57 | 0.874 | 0.99 | 0.53 | 0.64 | 3.60e-3 | 1.27 | 3.86e-3 | 0.50 | 1.00 | -0.02 | 0.80 |
| ENSG00000156976 | 1.00 | 8.15e-167 | 0.34 | 2.70e-17 | 0.59 | 0.70 | 0.43 | 0.62 | 0.56 | 0.65 | 0.15 | 0.75 | 0.74 | 2.48e-244 | 1,21 | 9.51e-301 |

### Supplementary Figure 5

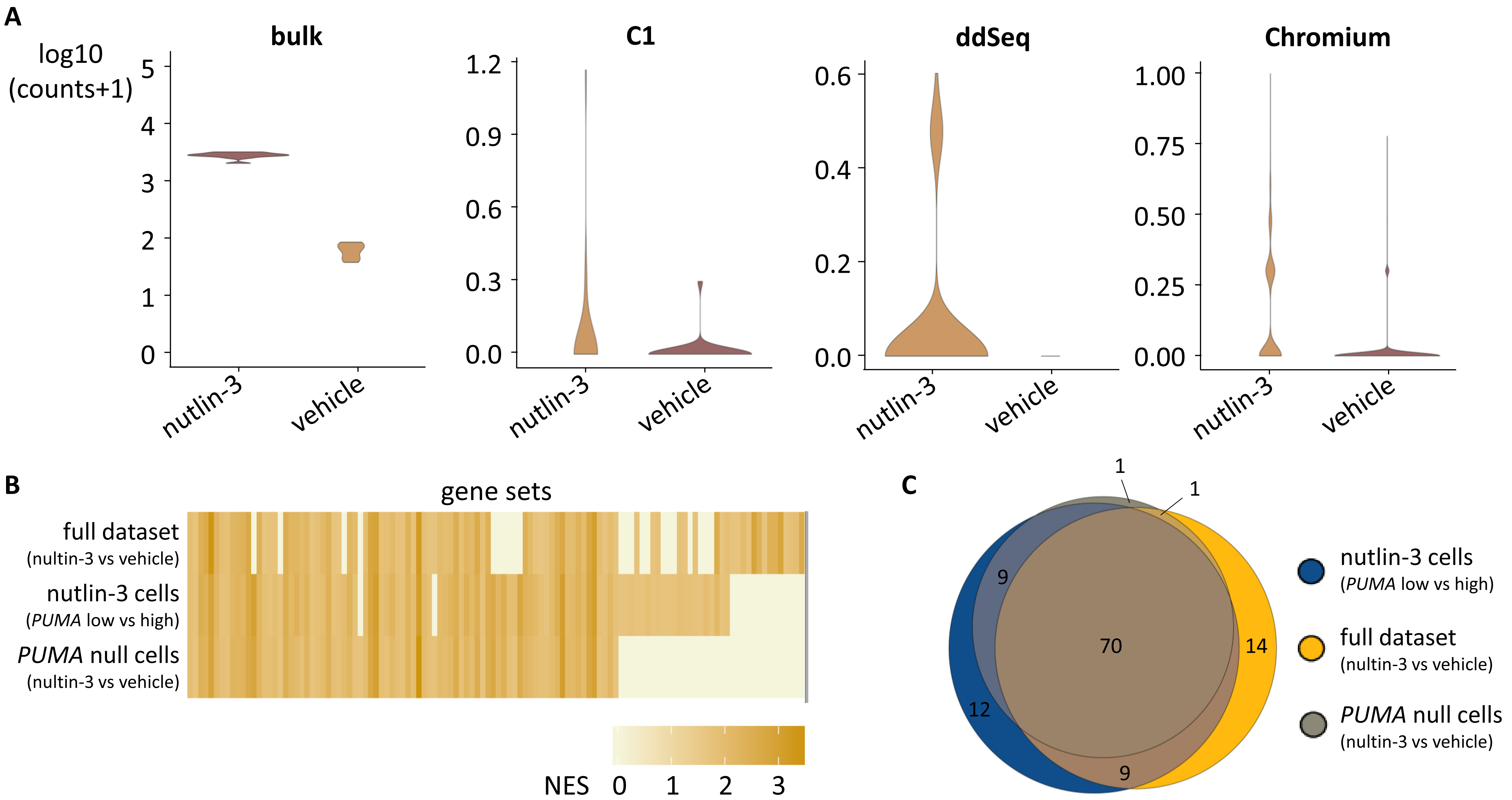

### Supplementary Figure 6

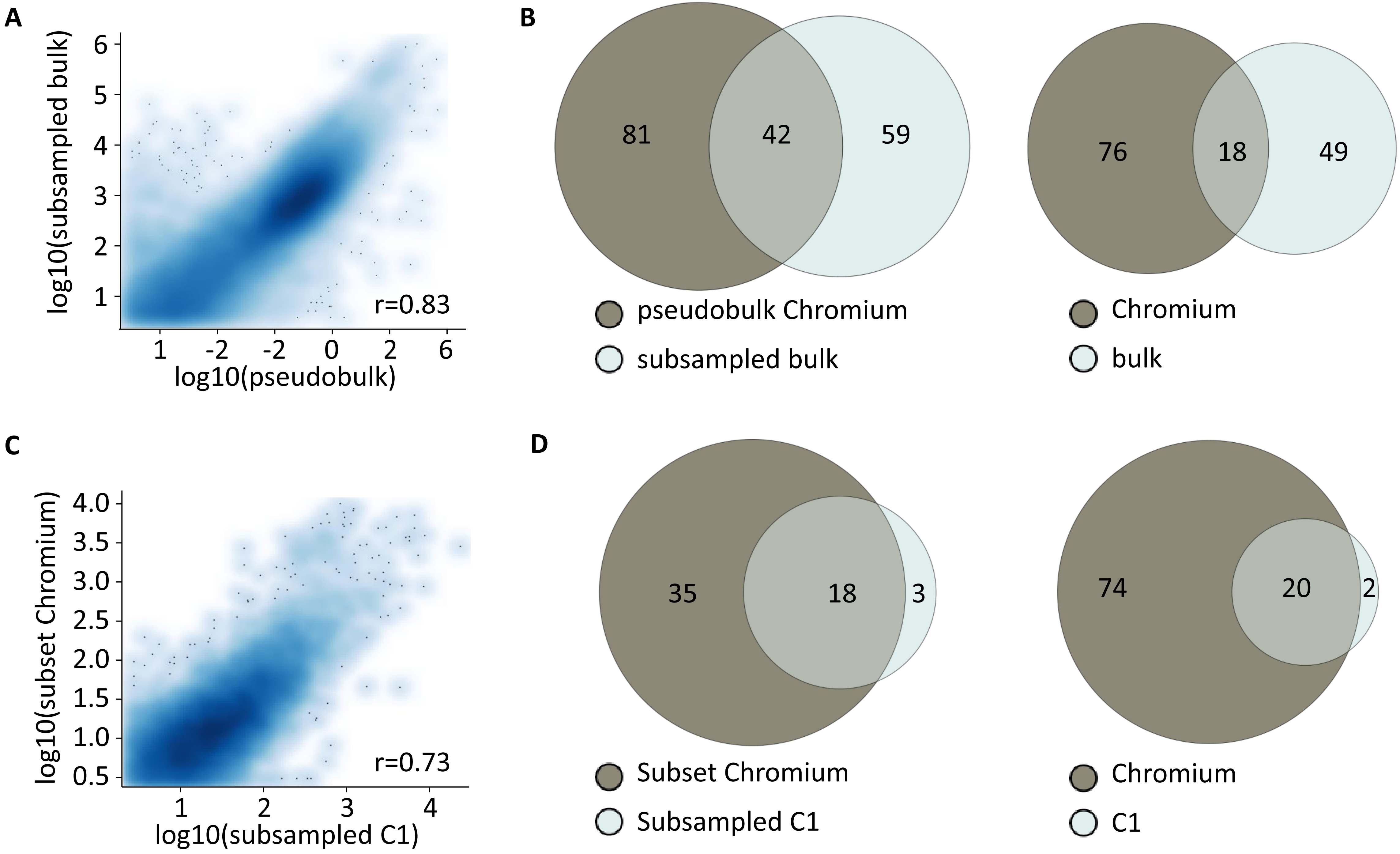

### Supplementary Figure 7

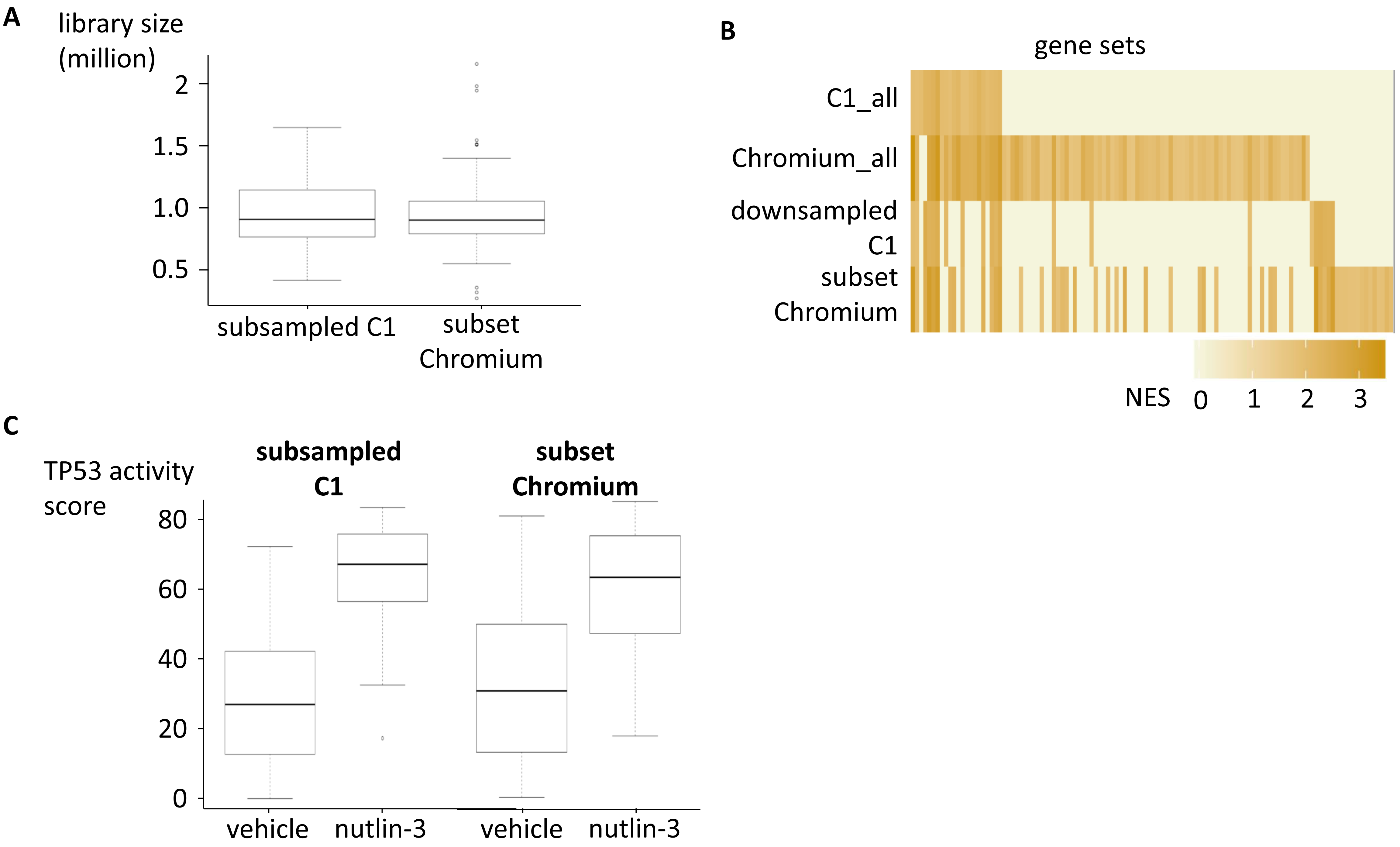

**Supplementary Figure legends**

**Supplementary Figure 1: cell cycles profiles and technical validation of the single cell RNA libraries.** (A) Representative example of a cell cycle profile of parental cells (left) and nutlin-3 treated cells (right). (B) RT-qPCR validation of TP53 target gene *CDKN1A* expression levels. Barplot shows the mean expression for the three single cell experiments. Bars represent the standard error of measurement (SEM). (C) Fragment Analyzer profile for C1 RNA libraries (left), Bioanalyzer profiles for ddSeq (middle) and Chromium (right).

**Supplementary Figure 2: gene expression correlation plots show high correlation between the devices.** (A) Smoothscatter plots showing the correlation in gene expression between C1, ddSeq and Chromium. (B) Smoothscatter plots showing the correlation in gene expression between bulk and each of the single cell devices.

**Supplementary Figure 3: quality assessment of the single cell RNA-seq data.** (A) PCA plots show a separation between nutlin-3 and vehicle treated cells for all devices. (B) Percentage of counts mapping on the top 25 highest expressed genes for the three single cell devices. (C) Overlap of the top 25 expressed genes for the three single cell devices.

**Supplementary Figure 4: violin plots for significantly differentially expressed genes in all or only one of the datasets.** (A) Expression of ENSG00000124762 (*CDKN1A*), significantly differentially expressed in all datasets. (B) Expression of ENSG00000004700 (*RECQL*), uniquely significantly differentially expressed in the bulk data. (C) Expression of ENSG00000160752 (*FDPS*), uniquely significantly differentially expressed in the C1 data. (D) Expression of ENSG00000179456 (*ZBTB18*), uniquely significantly differentially expressed in the ddSeq dataset. (E) Expression of ENSG00000156976 (*EIF4A2*), uniquely significantly differentially expressed in the Chromium dataset.

**Supplementary Figure 5: differences between cells with varying *PUMA* expression.** (A) *PUMA* expression in the bulk and single cell RNA-seq datasets. (B) Heatmap of significantly positively (q-value <0.05) enriched gene sets after GSEA for the C2-curated gene sets for each subgroup. Gene sets are color-coded according to their normalized enrichment score (NES). (C) Overlap between significantly positively enriched gene sets for the two cellular subgroups and full Chromium dataset.

**Supplementary Figure 6: pseudobulk data resembles true bulk data.** (A) Smoothscatter plot showing the correlation in gene expression between the pseudobulk and downsampled bulk dataset. (B) Overlap of significantly positively (q-value < 0.05) gene sets in the pseudobulk and downsampled bulk dataset and the original bulk and Chromium dataset. (C) Smoothscatter plot showing the correlation in gene expression between the downsampled C1 and subset of the Chromium dataset. (D) Overlap of significantly positively (q-value < 0.05) gene sets in the subsampled C1 and subset of the Chromium dataset.

**Supplementary Figure 7: comparing Chromium and C1 dataset for an equal number of cells at equal sequencing depth.** (A) After downsampling of the C1 data and taking an equal number of cells for the Chromium and C1 datasets, the mean number of reads per sample is 0.8 million. (B) Heatmap of significantly positively (q-value <0.05) enriched gene sets after GSEA for the C2 curated gene sets for the original, downsampled C1 and subset of Chromium datasets. (C) Boxplots depicting the TP53 activity score per cell, whereby ranking was based on the expression of 116 TP53 target genes.

**Supplementary Table 1. RT-qPCR primer sequences**

| | |
|---|---|
| CDKN1A_F | CCTCATCCCGTGTTCTCCTTT |
| CDKN1A_R | GTACCACCCAGCGGACAAGT |
| BAX_F | GATGCGTCCACCAAGAAGCT |
| BAX_R | CGGCCCCAGTTGAAGTTG |
| BBC3_F | CCTGGAGGGTCCTGTACAATCT |
| BBC3_R | GCACCTAATTGGGCTCCATCT |
| SDHA_F | TGGGAACAAGAGGGCATCTG |
| SDHA_R | CCACCACTGCATCAAATTCATG |
| TBP_F | CACGAACCACGGCACTGATT |
| TBP_R | TTTTCTTGCTGCCAGTCTGGAC |
| YWHAZ_F | ACTTTTGGTACATTGTGGCTTCAA |
| YWHAZ_R | CCGCCAGGACAAACCAGTAT |
| HPRT1_F | TGACACTGGCAAAACAATGCA |
| HPRT1_R | GGTCCTTTTCACCAGCAAGCT |

**Supplementary Table 2: number of significantly differentially expressed genes determined by EdgeR and PIM and number of positively enriched genesets.**

EdgeR is used in combination with Zinger for the single cell RNA sequencing datasets.

| | bulk | C1 |
|---|---|---|
| EdgeR (FDR < 0.05, abs(LFC) > 1) | 8303 | 52 |
| PIM (p.adj < 0.05, PI > 0.6 or PI < 0.4) | 17327 | 301 |
| EdgeR+PIM | 7010 | 40 |
| positively enriched genesets (q < 0.05) | 67 | 22 |

| | ddSeq | Chromium |
|---|---|---|
| EdgeR (FDR < 0.05, abs(LFC) > 1) | 107 | 97 |
| PIM (p.adj < 0.05, PI > 0.6 or PI < 0.4) | 193 | 502 |
| EdgeR+PIM | 28 | 88 |
| positively enriched genesets (q < 0.05) | 39 | 94 |

| | low_high_PUMA | G1 |
|---|---|---|
| EdgeR (FDR < 0.05, abs(LFC) > 1) | 106 | 112 |
| PIM (p.adj < 0.05, PI > 0.6 or PI < 0.4) | 494 | 504 |
| EdgeR+PIM | 86 | 105 |
| positively enriched genesets (q < 0.05) | 100 | 100 |

| | zero_CDKN1A | zero_PUMA |
|---|---|---|
| EdgeR (FDR < 0.05, abs(LFC) > 1) | 88 | 87 |
| PIM (p.adj < 0.05, PI > 0.6 or PI < 0.4) | 442 | 459 |
| EdgeR+PIM | 71 | 72 |
| positively enriched genesets (q < 0.05) | 73 | 81 |

| | bulk_subsampled_to_Chromium | Chromium_pseudobulk |
|---|---|---|
| EdgeR (FDR < 0.05, abs(LFC) > 1) | 5765 | 1361 |
| PIM (p.adj < 0.05, PI > 0.6 or PI < 0.4) | 12163 | 6358 |
| EdgeR+PIM | 4606 | 810 |
| positively enriched genesets (q < 0.05) | 101 | 123 |

| | subset_of_Chromium | subsampled_C1 |
|---|---|---|
| EdgeR (FDR < 0.05, abs(LFC) > 1) | 96 | 131 |
| PIM (p.adj < 0.05, PI > 0.6 or PI < 0.4) | 259 | 141 |
| EdgeR+PIM | 79 | 71 |
| positively enriched genesets (q < 0.05) | 53 | 21 |
